# Divergent habitat selection across many loci maintains species boundaries during introgression

**DOI:** 10.64898/2026.05.28.728564

**Authors:** Julia G. Harenčár, Kathryn Uckele, Kathleen M. Kay

## Abstract

Evidence of introgression between well-defined species is abundant, begging the question, how are species boundaries maintained during introgression? Work on this question has largely focused on systems where divergence is controlled by a few large effect genetic loci. However, adaptive divergence is often highly polygenic, especially between young species that remain capable of gene exchange. Here, we use phylogeny based introgression statistics and local ancestry inference to characterize the genomic, spatial, and historical extent of introgression between two recently diverged Neotropical plant species. We then use QTL analysis to investigate the genetic basis of traits under divergent abiotic and biotic selection that are involved in habitat isolation. Finally, we combine these top-down and bottom-up approaches to clarify how species with polygenic reproductive isolation maintain cohesion in the face of gene flow. We find clear evidence of both recent and ancient introgression between *Costus villosissimus* and *C. allenii*, but overall genomic divergence remains relatively high (*F*_*st*_:∼ 0.27) in part due to strong divergent habitat selection. Traits involved in divergent habitat adaptation are polygenic, such that strong habitat selection is spread across many loci rather than concentrated on a few loci of large effect. In contrast to the ‘islands of divergence’ seen around large effect loci in other hybridizing species pairs, we see genomically widespread and moderate peaks of both differentiation and introgression. Our results indicate that strong selection spread across many loci contributing to reproductive isolation can maintain species differentiation despite introgression.

**Significance Statement:** Gene flow between species (introgression) reduces genetic differentiation. And yet, introgression between well-defined species is common. How do species remain distinct in the face of introgression? Previous work focuses on differentiation maintained by elevated divergence in a few genomic regions (loci) with large effects. However, many loci with small effects commonly control differences between species. To clarify how species with abundant differentiating loci can remain distinct during introgression, we described patterns of introgression between two Neotropical plants and characterized the genetic basis of their divergent habitat adaptation. We found that isolation by divergent habitat adaptation is likely controlled by numerous small-effect and genomically widespread loci, and that habitat selection across multiple traits is sufficient to maintain differentiation despite ongoing introgression.

## Introduction

Gene flow between species is prevalent across the tree of life (Hedrick 2013; Harrison and Larson 2014; Mallet *et al*. 2016; Aguillon *et al*. 2022; Schley *et al*. 2022; Stull *et al*. 2023). This introgression can reverse divergence and break down isolating barriers (Grant 1981; Todesco *et al*. 2016; Ostevik *et al*. 2021). However, abundant evidence shows that species often remain distinct despite introgression (Whitney *et al*. 2010; Hedrick 2013; Harrison and Larson 2014; Aguillon *et al*. 2022; Schley *et al*. 2022; Stull *et al*. 2023). Most studies to date focus on cases where strong selection against gene flow is concentrated on a traits controlled by few large-effect genomic regions (loci) (Feder *et al*. 2012; Martin *et al*. 2013; Poelstra *et al*. 2014; Malinsky *et al*. 2015; Pearse *et al*. 2019; Wessinger *et al*. 2023). However, isolating traits are often controlled by many, rather than few, loci. Such polygenic divergence is common, especially for species isolated by complex traits like behavior and habitat adaptation (Fishman *et al*. 2002; Arnegard *et al*. 2014; Porth *et al*. 2015; Haasl and Payseur 2016; Exposito-Alonso *et al*. 2018; Rose *et al*. 2018; Martin *et al*. 2019; Barghi *et al*. 2020; Fagny and Austerlitz 2021; Huang *et al*. 2012b; He *et al*. 2023; Lazic *et al*. 2024; Nyine *et al*. 2025; Fifer *et al*. 2026). In these common situations, selection against introgression will be spread across the genome, potentially creating stable genome-wide divergence (Kautt *et al*. 2020) rather than genomic ‘islands’ of divergence (Turner *et al*. 2005; Turner and Hahn 2010). More generally, we expect different genomic underpinnings of species cohesion when many versus few loci underlie species divergence (Butlin and Smadja 2018; Merrill *et al*. 2024). Therefore, prevailing frameworks of localized genomic divergence may overlook the genome-wide mechanisms most relevant to complex adaptive divergence.

Multiple traits often interact to generate reproductive isolation, particularly in the early stages of speciation. Early stages of divergence tend to be dominated by prezygotic isolation, from divergence in behavior or habitat adaptation (reviewed in: Christie *et al*. 2022; Merrill *et al*. 2024). These early acting barriers are also frequently incomplete, allowing some gene flow between young species. This pattern is characteristic of young, rapid species radiations, where numerous closely related species co-occur across restricted geographic regions and experience some gene flow, but are largely isolated by divergent habitat adaptation (Jaramillo *et al*. 2008; Nicholls *et al*. 2015; Vargas *et al*. 2020; Schley *et al*. 2022; Zhao *et al*. 2024; Uckele *et al*. 2024). Rapid species radiations commonly occur alongside introgression, particularly when species are radiating into different ecological niches through complex trait divergence (Anderson 1949; Dowling and DeMarais 1993; Rieseberg *et al*. 1999; Beltrán *et al*. 2002; Salzburger *et al*. 2002; Smith *et al*. 2003; Seehausen 2004; Keller *et al*. 2013; Seehausen 2013; Meier *et al*. 2017; Kagawa and Takimoto 2018; Malinsky *et al*. 2018a; Meier *et al*. 2019; Svardal *et al*. 2020; Singhal *et al*. 2021; Schley *et al*. 2022; Zhao *et al*. 2024). The frequency and importance of complex habitat divergence in spite of - or even potentially facilitated by - introgression highlights the need to clarify the genomic underpinnings of polygenic reproductive isolation with gene flow.

Rapid species radiations and high rates of sympatry between close relatives are important contributors to biodiversity in the most species rich regions, like the tropics (Gentry 1982; Gentry *et al*. 1989; Pennington *et al*. 2015). For example, Neotropical plant clades boast diversification rates among the highest yet documented (Kay *et al*. 2005; Madriñán *et al*. 2013; Lagomarsino *et al*. 2016). Isolation due to divergent habitat adaptation is prevalent in these systems, as is hybridization, yet introgression is much less studied in tropical than temperate systems (Schley *et al*. 2022). How are so many close relatives co-occurring in these radiations despite common introgression? Some evidence suggests that genomically widespread, polygenic divergence is more effective at facilitating species divergence and cohesion during introgression than simple genetic architectures (Kautt *et al*. 2020; Svardal *et al*. 2020). However, we still lack understanding of the underlying interactions between introgression and the genetic basis of complex divergent adaptation to biotic and abiotic selection pressures.

Here, we address this gap in understanding by investigating whether the ‘islands’ of divergence framework applies when adaptive divergence involves multiple complex traits. We investigated two recently diverged tropical plant species with exceptionally well-characterized divergent habitat adaptation. Previous research in this system provides an understanding of the importance of habitat isolation and the traits involved that is rare for natural systems. *Costus allenii* (Jacq.) and *C. villosissimus* (Maas) are part of a diverse, recently and rapidly radiated clade of long-lived Neotropical understory plants, in which the species remain interfertile and are isolated by strong prezygotic barriers (Kay and Schemske 2008; Vargas *et al*. 2020; Kay and Grossenbacher 2022a; Uckele *et al*. 2024). *Costus allenii* and *C. villosissimus* cannot survive in each other’s habitat, as documented by multiple years of reciprocal transplant field experiments, and this is the primary isolating barrier between these otherwise interfertile, pollinator-sharing, and broadly sympatric species (Figure 2B; Chen 2011, 2013; Chen and Schemske 2015. *Costus allenii* inhabits perennially wet, dark forest understory with high herbivore pressure; accordingly, *C. allenii* has tough, well defended leaves and no seed dormancy, but grows relatively slowly. *Costus villosissimus* inhabits high-light forest edge habitat with seasonal drought and low herbivore pressure. It is effectively drought deciduous, actively senescing its thin, poorly-defended leaves in the dry season and rapidly regrowing them during the rainy season. Seeds are produced at the end of the wet season and lie dormant through the following dry season Chen 2011, 2013; Chen and Schemske 2015; Harenčár *et al*. 2022, 2024. Thus, a suite of coordinated traits facilitates adaptation to one habitat versus the other, with strong trade-offs between them. We first evaluated how effectively habitat isolation limits genetic exchange by characterizing recent and ancient hybridization and quantifying genomic patterns of introgression. Next, to clarify the genetic basis of multiple traits involved in divergent habitat adaptation (see Figure 1 for a description of divergent traits), we identified quantitative trait loci (QTL) with a greenhouse-grown genetic mapping population. Although we don’t yet know the relative strength of selection on each individual trait involved in habitat isolation, which would require infeasible large-scale field experiments with hybrids, we investigated key traits representing habitat isolation across several categories: physiology, life history, and defense. Finally, we compared genomic patterns of differentiation to the genetic basis of divergent habitat adaptation to understand how known ecological barriers impact the genetics of reproductive isolation. This integration of ecological understanding with top-down and bottom-up genomic approaches provides rare insight into how polygenic adaptive divergence can persist despite introgression.

**Figure 1.**
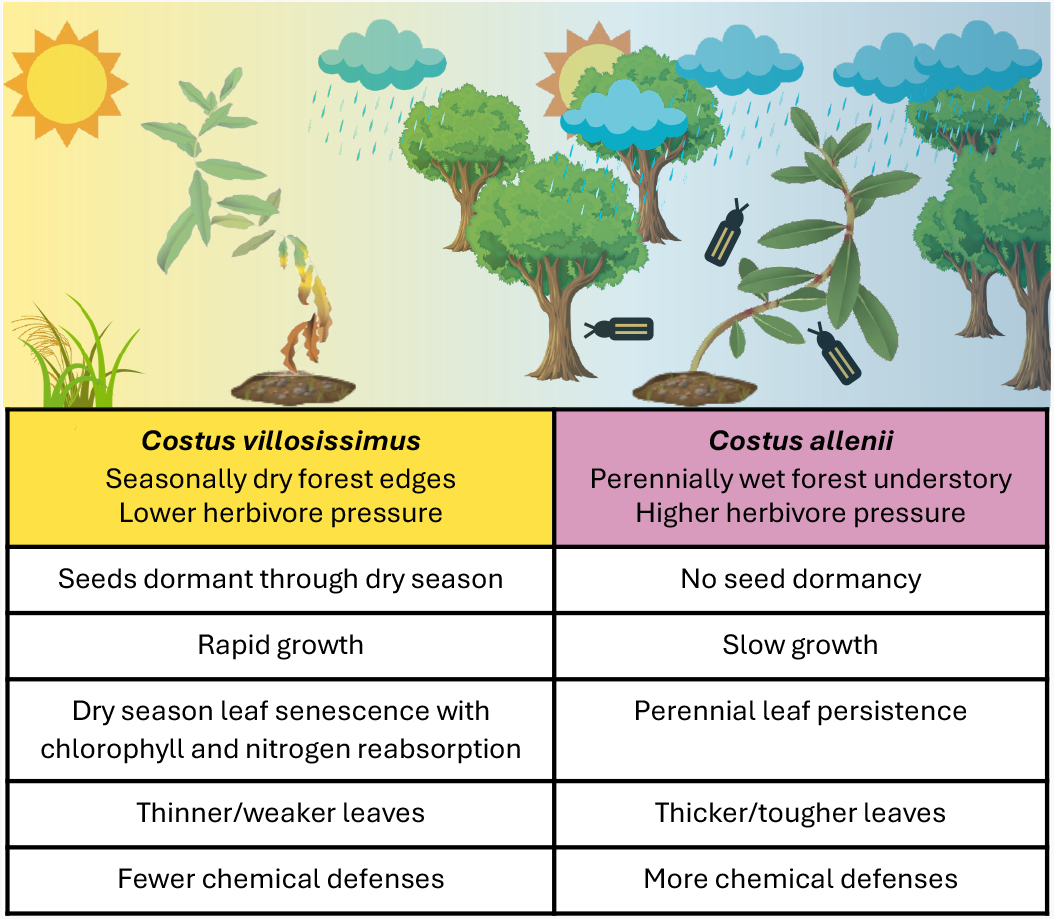
Habitat isolation of *Costus villosissimus* and C. *allenii*. The diagram illustrates typical habitat for wild C. *villosissimus* (left) and C. *allenii* (right); drier, higher light, and more open habitat for C. *villosissimus*, and wetter, lower light, closed canopy habitat with more herbivores for C. *allenii* (Chen 2011, 2013; Chen and Schemske 2015; Harenčár *et al.* 2022, 2024). We conducted QTL analysis for each trait in the table other than chemical defenses.

**Figure 2.**
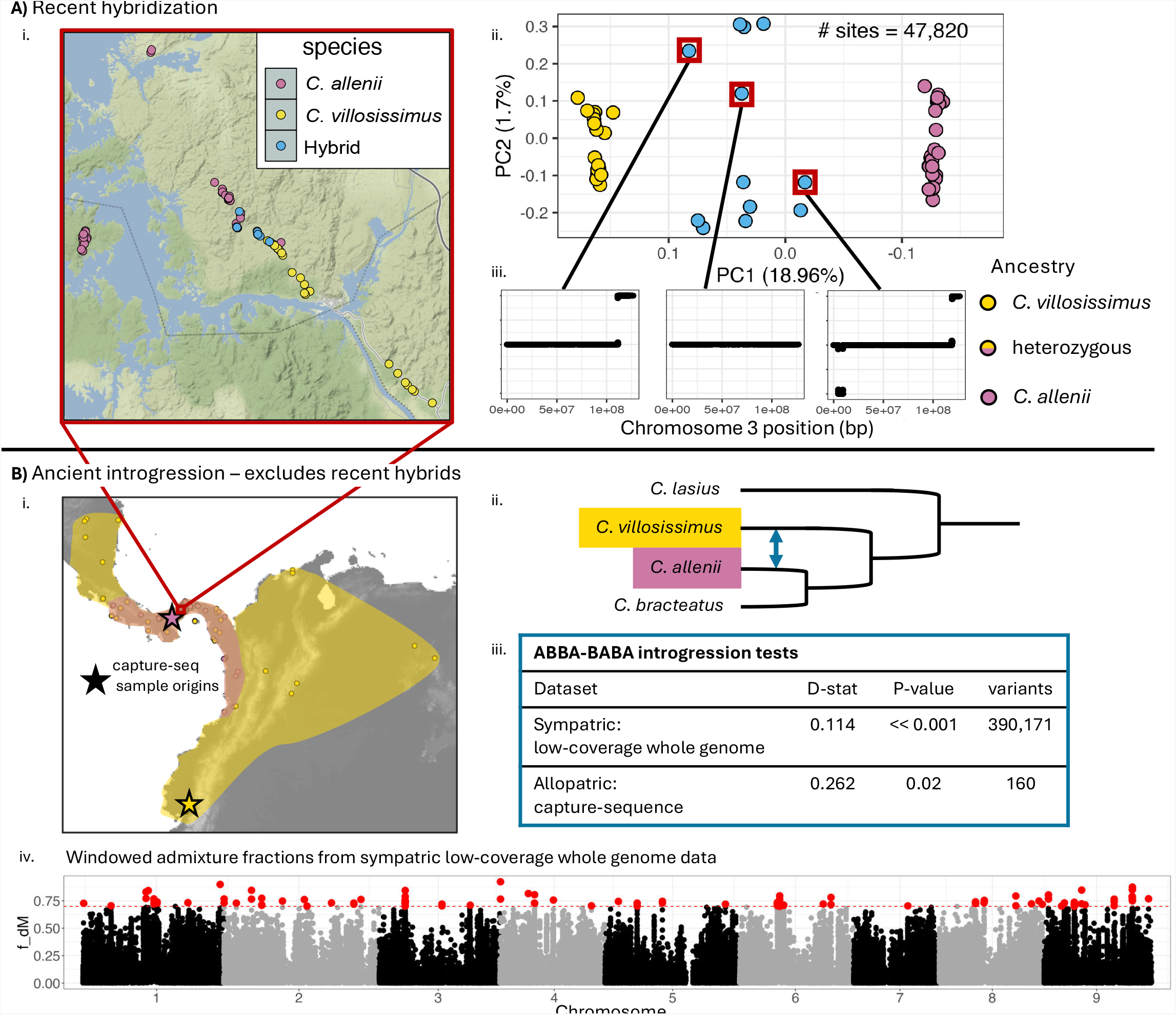
Recent and ancient introgression. **A)** Evidence of recent hybridization. i. A map of population genetic samples from Panama *colored* by species. ii. A genetic PCA built from 47,820 SNPs that are shared across 80% of individuals of each taxon. iii. Example local ancestry plots for chromosome 3 of the individuals boxed in red on the PCA. The x-axis is position along the chromosome and the y-axis represents ancestry at that position, with 0 indicating homozygous C. *allenii* ancestry, 0.5 indicating heterozygous ancestry, and 1 indicating homozygous C. *villosissimus* ancestry. From left to right, individuals are examples of a backcross to C. *villosissimus*, an F1 hybrid, and a F2 or further generation hybrid. **B)** Evidence of ancient introgression. i. Map depicting C. *allenii*’s range in pink and C. *villosissimus*’s range in yellow. Circles show confirmed occurrences and stars indicate the individuals with capture sequencing data used to test for allopatric evidence of introgression. ii. Phylogeny of the focal clade with in- and out-group species used for ABBA-BABA tests. iii. Whole genome introgression analysis results from both the population genetic low-coverage whole-genome data (not including hybrids from panel A) and from the capture sequence data from allopatric individuals. iv. Genomic patterns of introgression. The x-axis is position along each chromosome and the y-axis is f_*dM*_ calculated in 50 variant sliding windows of 25 variant steps. The red dashed line represents the 99.9th percentile of negative values, and red dots represent genomic windows with evidence of introgression.

## Results

### Recent and ancient introgression

#### Hybrids persist in intermediate habitat

We collected and sequenced samples of both species and putative hybrids from Panama, where the species co-occur (Figure 2A). Recent hybridization (within 10s of generations) between *C. allenii* and *C. villosissimus* was evident from both PCA and local ancestry inference, which identifies tracks of heterozygous and homozygous parental ancestry in recent hybrids. The PCA clearly separated species on PC1 (18.96% variance explained), revealing 12 *C. allenii* - *C. villosissimus* hybrids, which were all found in habitat with intermediate light and precipitation near the center of the Panamanian isthmus (Figure 2A). Local ancestry inference analysis revealed that five of the 12 hybrids are F1s, three are likely backcrosses to *C. villosissimus* and four are F2 or later-generation hybrids (Figure 2A and Figure S1). Even the potential further generation hybrids have large ancestry blocks indicating only a few generations of hybridization. Given potentially long generation times in *Costus*, the recency of this hybridization suggests it may be facilitated by anthropogenic disturbance; we collected these hybrids along Pipeline Road, which was built during World War II (early 1940’s) as an alternate route for oil transport. The road creates a light gap that cuts across the precipitation gradient of the Panamanian isthmus, providing a pathway for *C. villosissimus* to come in close contact with *C. allenii* near the center of the precipitation gradient. These relatively recent hybrids were excluded from all further analyses as they are not informative for investigating genomic patterns of historical introgression.

#### Historical introgression is widespread across the landscape and genome

While Pipeline Road likely increases the chances of hybridization, the two species’ habitats can naturally occur within the dispersal distance of their orchid bee pollinators (Janzen 1971), and potentially within bird dispersal distance of their seeds (though bird dispersal has not been confirmed). To quantify historical introgression (which predates recent anthropogenic disturbance in Panama), we excluded all individuals identified as hybrids in the PCA and local ancestry inference and used the remaining ‘pure’ individuals to run analyses designed to identify signals of ancient introgression. We ran two separate ABBA-BABA analyses (Figure 2A.i). In the first, we ran the analysis on each chromosome and the whole genome using low-coverage sequencing of the Panamanian samples of each species, which are from the sympatric portion of the species range (Figure 2B.iii). We detected strong evidence for introgression on seven of nine chromosomes, marginal evidence on one, and one without a significant overall D-statistic (Table S1). Across all chromosomes, including those with marginal or non-significant D-statistics, two sliding window estimates of introgression signal (*d* _*f*_ and 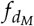) revealed consistent regions of significant introgression (Figures 2B.iv and S2). We ran the second ABBA-BABA analysis with previously published capture-sequence data from a Panamanian *C. allenii* individual and a *C. villosissimus* individual from outside *C. allenii*’s range (allopatric comparison; Vargas *et al*. 2020). This analysis had fewer loci (25,987 SNPs from within 756 genes) and thus was only run at the whole genome scale, again revealing significant introgression and indicating that the signal of introgression is range-wide and not limited to the *C. allenii* and *C. villosissimus* in Panama (Figure 2B.iii and Table S1).

### Quantitative trait loci explain limited trait variance

Using a mapping population of 390 F3s, we identified nine QTL across five of the six traits involved in divergent habitat adaptation (Table S2). Model fitting revealed no QTL for leaf thickness. QTL were distributed across five of the nine chromosomes and only two co-localized (Figure S4). Each QTL explained relatively small amounts of the phenotypic variance, ranging from 1.2%-8.4% (except for a seed dormancy QTL, which explained 17.8% of variance; Figure 3A and Table S2). For seed dormancy, the two QTL identified explained 26.2% of trait variance. For the other traits, QTL only explained 3-8% of trait variance. In all these cases, genetic background (the F2 family F3s were from and/or hybrid index) and environment (greenhouse cohort) each explained more trait variance than the QTL, with variance explained by cohort ranging from 21.9-28.9% and by family from 5.0-27.1%. The total unexplained variance was high across all QTL models, ranging from 37-64% (Figure 3A). Unexplained variance is caused by either phenotypic plasticity due to environmental variation, or additional genetic loci with sub-threshold effects on the trait. Here, we captured the majority of environmental variability experienced by experimental mapping population plants with the cohort variable; plants in the same cohort moved through growth chambers and greenhouse benches together and were never more than roughly a meter apart. Thus, we infer that a substantial portion of trait variance was due to additional genetic loci of low effect. Additional loci of low effect were also suggested by trait association genome scans showing numerous unevenly distributed sub-threshold peaks (Figures 3B and S3). Further supporting the likelihood of additional undetected loci, we found several QTL with effects in the opposite direction expected based on parental values (antagonistic QTL). One of the two QTL for seed dormancy and growth rate as well as both toughness QTL were antagonistic (Table S2). Interestingly, while the QTL for both nitrogen and chlorophyll reabsorption during drought were antagonistic in the model with covariates (including hybrid index and F2 family), the raw genotype means at those loci were in the opposite and expected direction (Table S2). This sign flip when genetic background is accounted for with covariates indicates that there is only an antagonistic effect at these QTL if we hold genetic background as constant. In other words, homozygous ancestry for the more drought resistant parent at those specific QTL is associated with genome-wide ancestry for that parent species; the association between the genotype at that locus and genome-wide ancestry is likely why the raw QTL genotype means associate as expected with nutrient reabsorbance during drought. The association of genome-wide ancestry with these QTL and their effects thus strongly supports a polygenic basis for these important drought adaptation traits.

**Figure 3.**
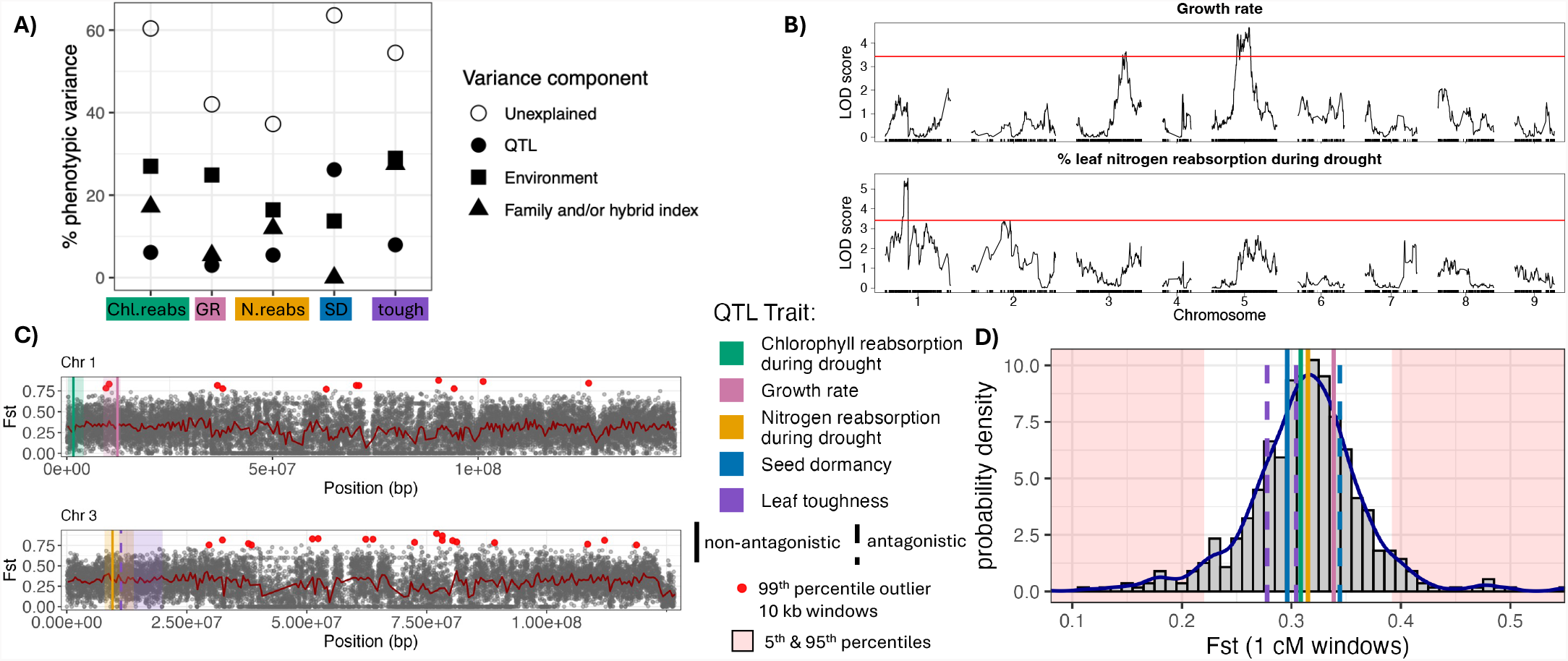
QTL explain little trait variance and do not have elevated divergence. **A)** Percent of trait variance explained by QTL (filled circles); the plant cohort, which captures most of the environmental variability (squares); the plant family defined by F2 parent and/or hybrid index, depending on which were included by model selection (triangles); and the trait variance not explained by the model (open circles). **B)** Example QTL peaks with 10% LOD score cutoffs in red calculated from 1000 permutations of scanone indicating numerous subthreshold peaks involved in polygenic control of the traits. **C)** Genome wide F_*st*_ calculated in 10 kb windows with overlaid QTL. Red points are 10 kb windows in the 99.9th percentile and the red line indicates F_*st*_average calculated in 500 kb windows. Colored bars indicate QTL peaks with dashed bars indicating antagonistic QTL and shaded regions showing Bayes 95% credible intervals (CIs). Two representative chromosomes are shown here, for all chromosomes see Figure S4. **D)** Average F_*st*_ in the seven QTL CIs smaller than 15 cMs compared to genome-wide F_*st*_ averages. Dashed QTL bars again indicate the antagonistic QTL, grey bars display the frequency of values averaged for non-overlapping 1 cM windows, and red shading indicates the 5th and 95th percentiles. QTL CIs range from 0.85-14.8 cMs (Table S2).

### QTL associated with habitat isolation do not show consistently elevated divergence

Despite widespread introgression and strong divergent adaptation between species, we did not see a patten of consistently elevated introgression at antagonistic QTL, or elevated divergence at non-antagonistic QTL involved in habitat isolation. Overall divergence between *C. allenii* and *C. villosissimus* calculated as Weir & Cockerham’s *F*_*st*_ (1984) with variance components summed across 50 kb non-overlapping windows across the genome was 0.27. Local patterns of differentiation across the genome displayed widespread peaks of divergence, with various outlier *F*_*st*_ 10 kb windows across all chromosomes (Figures 3, S2, and S4). However, the seven reasonably sized QTL credible intervals (CIs; two were removed because of spanning over 100 Mbp, the rest span less than ∼10 Mbp/15 cMs) did not have reduced or elevated divergence (or introgression) relative to genome-wide averages (Figures 3 and S5). Some signal may be obscured by large CIs (plotted CIs range from 0.85 to 15 cM), but only two of seven QTL CIs contained *F*_*st*_ outlier windows (nitrogen reabsorption on chromosome one and leaf toughness on chromosome eight; Figure S4).

## Discussion

Introgression is common between lineages that remain distinct. This contradictory outcome is best understood in systems where few large-effect loci control reproductive isolation, and remain distinct despite introgression (i.e. ‘Islands’ of divergence; Turner *et al*. 2005; Turner and Hahn 2010). However, reproductive isolation is polygenic for many species, especially in cases of habitat isolation, where many adaptive traits work in concert. Habitat isolation is common in recently diverged species, which often lack postzygotic barriers and can hybridize. Here, we leverage the comprehensive characterization of various traits involved in divergent habitat adaptation between *C. allenii* and *C. villosissimus* to clarify the genetic basis of habitat isolation and relate it to genome-wide patterns of introgression. We found that the genetic basis of divergent habitat adaptation is polygenic, and signals of both introgression and differentiation are spread across the genome rather than restricted to a few regions. Detectable QTL explain only small portions of trait variance, and do not show the elevated divergence seen in similar studies on few large effect QTL (Renaut *et al*. 2013; Pearse *et al*. 2019; Todesco *et al*. 2020; Coughlan *et al*. 2021; Powell *et al*. 2021; Wessinger *et al*. 2023; Schield *et al*. 2024). Our results suggest that when reproductive isolation is highly polygenic, selection on any one locus is weak and even loci contributing to important differentiating traits will not be entirely resistant to introgression. Rather, selection on polygenic traits creates diffuse, genome-wide selection that limits introgression and maintains overall genomic divergence without strong signals at specific loci (supporting theory in Veller *et al*. (2023); Veller and Simons (????); Ragsdale (2025); Negm and Veller (2026)).

The genetic basis of isolation between our focal species is not localized to a few genomic regions. Adaptively divergent trait QTL only explained a small portion of trait variance, whereas overall genetic background (captured by hybrid index and/or family representing additional genetic effects) generally explained more variance than the QTL. Further, there were large portions of unexplained variance (37%-64%), consistent with additional genetic effects not captured by QTL (Lynch *et al*. 1998; Broman *et al*. 2009). Additional undetected small effect loci are also likely because of relatively low power for QTL mapping with <400 individuals; much larger mapping populations are necessary to accurately detect low effect loci (Otto and Jones 2000; Beavis 2019). Even so, our genome scans displayed wide regions with modest LOD score elevation but a few sharp peaks that do not reach the LOD significance threshold, indicating many additional loci contribute to trait variance (Broman *et al*. 2009; Beavis 2019). We confirmed this interpretation by adding additional potential QTL to our simple models, conditional on the existing QTL; these analyses often revealed additional QTL with epistatic interactions with the original QTL, but are omitted here due to model complexity penalties. Polygenicity is further supported by the detection of antagonistic QTL. Antagonistic QTL are common and their primary explanation is that parental species carry many complementary additive alleles, such that each parent carries some alleles pushing the trait in the contrary direction to the overall phenotype, enabling a smooth distribution of trait values (Rieseberg *et al*. 1999, 2003). Antagonistic QTL can also reflect sign epistasis, in which the direction of the allele’s effect depends on interaction with additional loci that fall below detection thresholds, as is suggested by our nutrient reabsorption QTL having opposite effect directions after accounting for genetic background with hybrid index and F2 family (Huang *et al*. 2012a; Mackay 2014). Taken together, these QTL analyses results support a polygenic basis of divergent habitat adaptation between our focal species, consistent with other studies on the genetic basis of complex habitat adaptation (Fishman *et al*. 2002; Huang *et al*. 2012b; Rockman 2012; Arnegard *et al*. 2014; Porth *et al*. 2015; Haasl and Payseur 2016; Exposito-Alonso *et al*. 2018; Rose *et al*. 2018; Martin *et al*. 2019; Barghi *et al*. 2020; Fagny and Austerlitz 2021; He *et al*. 2023; Lazic *et al*. 2024; Nyine *et al*. 2025; Fifer *et al*. 2026).

Even though polygenic divergence frequently undergirds reproductive isolation, the majority of research on species divergence with introgression focuses on cases where a few moderate to large effect loci account for most of the variance in the focal reproductive isolating traits. Some recent work on migration-selection balance theory highlights that concentrated or simple genetic architectures of reproductive isolating loci facilitate species differentiation despite introgression (Bürger and Akerman 2011; Yeaman and Whitlock 2011; Yeaman 2022). Numerous empirical examples support this, often finding signals of divergence localized to relatively few large effect loci (Renaut *et al*. 2013; Pearse *et al*. 2019; Todesco *et al*. 2020; Coughlan *et al*. 2021; Powell *et al*. 2021; Wessinger *et al*. 2023; Schield *et al*. 2024). In studies exploring a greater number of moderate and low effect loci, the correlation between QTL for isolating traits and elevated genetic divergence tends to be weaker, with some larger effect loci showing elevated divergence, but not most QTL (Roda *et al*. 2017; Price *et al*. 2018). A few recent studies found the same lack of elevated divergence at reproductive isolation loci that we saw here. Frayer and Payseur (2024) collated well-characterized loci tied to postzygotic barriers in house mice species and found they do not coincide with loci where divergence is more elevated than expected by chance. Similarly, Stankowski *et al*. (2023) found no clear overlap between even large effect pollination syndrome QTL and barrier loci identified through population genomic scans. These examples fit well with recent theoretical work showing that when traits are highly polygenic, even very small amounts of migration lead to variation in the genetic architecture of local adaptation (Sakamoto *et al*. 2024), and that stabilizing selection can maintain trait divergence without high divergence at specific loci (Yair and Coop 2022). They also fit with classical migration-selection balance theory, which clarifies that polygenic divergence can withstand gene flow so long as migration is relatively low and selection is sufficiently strong (Haldane 1927; Wright 1931; Slatkin 1973; Nosil 2012; Flaxman *et al*. 2014; Barton and Etheridge 2018; Yeaman 2022). Strong selection may not even be necessary; recent theory demonstrates that even weak stabilizing selection on a polygenic trait will act to purge minor-parent ancestry genome-wide after introgression without leaving a signature at any particular locus (Veller and Simons ????), predicting exactly the empirical pattern we described. It is worth noting that this pattern stands even though we likely haven’t captured all traits important for habitat isolation; both *F*_*st*_ and introgression outliers are relatively evenly distributed across the genome rather than concentrated in a few dramatic peaks (Figures 2B and S2). Indeed, additional isolating adaptations would mean more loci and thus greater polygenicity of isolation. Polygenic architectures, rather than inhibiting divergence, may even promote stable and rapid sympatric speciation under divergent selection (Kautt *et al*. 2020; Svardal *et al*. 2020).

Habitat differences can create strong divergent selection; in our system, neither species can survive in the other’s habitat (Chen 2013; Chen and Schemske 2015). This strong selection maintains differentiation across the genome, with *F*_*st*_ in line with other recently diverged but clearly distinct species with detectable gene-flow (e.g. Strasburg *et al*. 2009; Martin *et al*. 2013). Even with this clear ecological and genetic divergence, there is genomically widespread signal of introgression (Figures 2B, Figure S2, and Table S1). The introgression peaks spread across the genome may represent relatively neutral loci, or loci experiencing positive selection on introgression (adaptive introgression) despite strong overall selection against introgression (Short and Streisfeld 2025). When complex traits like habitat adaptation are controlled by many loci, selection maintaining divergence is spread across those loci, such that any single locus experiences lowered selection against introgression.

Selection against introgression due to habitat also varies through time and space along with environmental variation, such that even moderate to large effect loci may not have clearly elevated divergence. As an example, consider the relatively larger effect (17.8% trait variance explained) antagonistic QTL for seed dormancy. Differences in seed dormancy are crucial for habitat isolation between these species; a seedling will not survive if the seed germinates before or during the dry season in *C. villosissimus* habitat. However, the antagonistic QTL is at least partially responsible for a few *C. villosissimus* seeds in wet conditions germinating immediately (Figure S6). Immediate germination would generally be fatal in the wild, but not always. Rare years with little or no dry season, or forest edges along lakes or rivers, might generate conditions that select for early germination in *C. villosissimus*. However, a *C. allenii* individual would still be maladapted in that environment (e.g., due to higher light and future drought events) such that there would still be selection against *C. allenii* ancestry. This example illustrates how strong but variable habitat selection acting on polygenic adaptations can allow for introgression at a particular locus while genome-wide selection against introgression enables species differentiation.

Like fluctuating selection, repeated hybridization can also obscure patterns of divergence at reproductive isolation associated loci. Both simulations and empirical evidence show that genomic patterns of differentiation vary substantially between replicate hybridization events (Ferchaud and Hansen 2016; McFarlane *et al*. 2024). If the different hybridization events also occur in differing environments, genomic patterns of introgression and differentiation will be even less predictable as the loci influencing a given polygenic trait are more likely to vary between populations (Lopez-Arboleda *et al*. 2021). Thus, gene flow following different hybridization events can mix distinct population level genomic patterns of divergence, precluding a clear pattern of divergence at specific reproductive isolation associated loci. Further, theory indicates that repeated, episodic gene flow can shrink the size of divergent loci (Flaxman *et al*. 2013), and that complex epistasis with many interacting divergent loci increases robustness to gene flow (Fraïsse *et al*. 2016; Kautt *et al*. 2020). In the context of this theory, our findings of diffuse, genome-wide divergence and introgression fit expectations given the interaction of a complex, polygenic basis of habitat divergence with strong but fluctuating selection pressures and repeated, occasional historical introgression.

In conclusion, introgression between *C. allenii* and *C. villosissimus* is widespread across space, time, and the genome, yet the species remain ecologically and genetically distinct. Unlike the majority of previous studies, we did not find genomic islands of divergence around a few key loci. Rather, strong habitat isolation is controlled by many loci of small to medium effect across spatially and temporally variable environments. Variable and genomically diffuse selection across loci permits introgression while preserving key differentiating adaptations. Given that both complex habitat isolation and introgression are common in adaptive radiations, our study provides an example framework for how diversification and introgression can coexist during adaptive radiations. Our study provides the first detailed empirical example of how polygenic divergent adaptation to both biotic and abiotic environmental conditions can maintain species distinctions despite ongoing introgression.

## Materials and Methods

### Study system

The Neotropical spiral gingers (*Costus*) form a large clade of Central and South American herbaceous perennial understory plants that have diversified into approximately 84 species within the last ∼ 3 million years (Vargas *et al*. 2020; Maas *et al*. 2023). These species generally lack postzygotic barriers, remaining broadly interfertile (Kay and Schemske 2008), but commonly display pre-mating barriers such as habitat and pollinator isolation (Kay and Schemske 2003; Kay *et al*. 2006; Chen 2011, 2013; Chen and Schemske 2015; Vargas *et al*. 2020; Kay and Grossenbacher 2022b). *Costus allenii* (Jacq.) and *C. villosissimus* (Maas) share pollinators and have overlapping ranges, but display strong habitat isolation (Figure 1; Chen 2011). This habitat isolation is the primary isolating barrier between *C. allenii* and *C. villosissimus* (Chen 2011, 2013; Chen and Schemske 2015). Chen and Schemske (2015) showed that *C. villosissimus* seedlings cannot survival in *C. allenii*’s perennially wet, dark forest understory habitat, and *C. allenii* cannot survive seasonal drought in *C. villosissimus*’s high-light forest edge habitat (Chen and Schemske 2015, 2019). In early life, *C. villosissimus* avoids drought through seed dormancy, germinating at the start of the wet season. As an adult, it achieves drought adaptation through flexible drought-deciduousness enabled by active leaf drop (senescence) in the dry season followed by rapid growth with the onset of the wet season (Chen and Schemske 2015; Harenčár *et al*. 2022). Rapid growth comes with a tradeoff in which *C. villosissimus* invests less in defense against herbivory, whereas *C. allenii* grows more slowly but has greater investment in chemical and structural defenses (Harenčár *et al*. 2024)). This lower defense investment in *C. villosissimus* aligns with differences in biotic selection pressures between habitats; the primary beetle herbivores of *Costus* require wet habitat, such that there is lower herbivore pressure in the seasonally dry, open habitat of *C. villosissimus* (Harenčár *et al*. 2024).

### Sample collection, DNA extraction, and sequencing

#### Wild sample collection

In Central Panama, *Costus allenii* and *C. villosissimus* are generally distributed along a precipitation gradient, with *C. villosissimus* primarily growing near forest edges on the seasonally dry southern side of the isthmus and *C. allenii* primarily growing in dark ravines on the perennially wet northern side of the isthmus (Figure 2A.i). Previous research noted potential wild hybrids near the center of the precipitation/seasonality gradient (Chen 2011). We collected samples from 38 *C. allenii* individuals, 23 *C. villosissimus* individuals, and, around the center of the seasonality gradient of precipitation, 12 hybrids. To facilitate tests for introgression, we also collected tissue from 21 individuals of *C. allenii*’s sister species, *C. bracteatus* (ingroup), from Costa Rica, and from two *C. lasius* (outgroup) individuals – one from Panama and one from Peru (Figure 2B.ii).

#### Genetic mapping population creation

We generated a mapping population of F3 hybrids by self-pollinating a wild F1 (identity confirmed through local ancestry assignment; see “Local ancestry assignment” section below), growing the resulting F2 seeds in the UCSC greenhouses, and self-pollinating and collecting seed from five of those F2s to generate 390 F3s. Producing an F3 mapping population in the greenhouse minimizes maternal effects from variable field conditions, while taking advantage of natural hybridization to provide an additional generation of recombination in these long-lived plants. We recorded the plant’s “cohort”, defined for seeds and seed dormancy as individuals sowed on the same day, and defined for all other measurements as plants receiving their first growth rate measurement and entering the growth chambers on the same day. These cohorts therefore capture most of the environmental variability experience by mapping population plants. This means that variance in a trait not captured by our QTL model is less representative of plasticity due to environment and more indicative of numerous contributing genetic interactions and loci of small effect. We collected and extracted DNA from the 390 F3s, the original wild F1 parent, and 21 F2s including the five F2 parents of the F3s.

#### DNA extraction, sequencing, and variant calling

For all DNA extractions, we used Macherey Nagel Plant II kits (Düren, Germany) on meristematic or flower bud tissue dried in silica. For the wild collected samples used in population genomic analyses, we generated libraries for low-coverage whole-genome sequencing with Tn5 transposase and tagmentation, then sent them to Fulgent Genetics for paired-end 150 bp read sequencing on an Illumina HiSeq4000. An additional 6 wild samples were sent together with all the mapping population individuals to the DNA Technologies Core at the UC Davis Genome Center for their High Throughput library preparation and sequencing (also Illumina 150pe).

We inspected FASTQ files with FastQC (Andrews 2010), trimmed adapters and low-quality bases with Trimmomatic v0.39 (Bolger *et al*. 2014), and removed PCR duplicates using the dedupe function in BBTools v39.01 (Bushnell 2014). Next, we mapped reads to the *C. lasius* genome (NCBI accession number GCA_027563935.1; Harenčár *et al*. 2023), using BWA mem version 0.7.15-r1140 (Li and Durbin 2009). Mapping to the *C. lasius* genome avoids reference bias, as *C. lasius* is sister to the small clade containing *C. allenii* and *C. villosissimus*, making it equally related to both focal species (Vargas *et al*. 2020). We removed ten *C. allenii* and three *C. villosissimus* from the wild collected samples with outlier low coverage (<0.15x) or low coverage breadth (<10%), and one outlier F3 sample with lower than 0.05X average coverage across the genome from future analyses. We filtered out unmapped reads, reads smaller than 35 bp, and reads with mapping quality lower than 20. We then calculated mappability with genmap (Pockrandt *et al*. 2020), using 150 bp kmers and a 100 bp max distance for merging regions, and filtered out regions with low mappability (below 0.7). When calling biallelic SNPs, we applied a base and map quality filter of 20 and a 0.05 MAF filter, removed sites within three base pairs of an indel, and removed sites with coverage depth greater than the 99th percentile.

### Population genetics of wild plants

#### Recent hybridization

We visualized patterns of gene flow between *C. allenii* and *C. villosissimus* with Principal Component Analysis (PCA). In addition to the filters described above, we increased base and mapping quality filters to a threshold of 30, and only included sites with at least 1x coverage in at least 80% of each species and the hybrids. We ran ANGSD (v gnu-937) to estimate genotype likelihoods with Samtools, using a 1e-6 p-value SNP calling cutoff, an excess heterozygosity cutoff of 1e-2, and inferring the major and minor alleles (Korneliussen *et al*. 2014). We then created the PCA from genotype likelihoods with `PCAngsd` .

#### Ancestry assignment across hybrid genomes

We mapped local ancestry across the genome for each individual that appeared to be a hybrid from the wild population genetic PCA using Ancestry HMM (Corbett-Detig and Nielsen 2017). We used sequences from 18 wild *C. villosissimus* and 24 wild *C. allenii* individuals that clearly separated to the ends of PC1 in the genetic PCA for the parent reference panels. When generating the aHMM input panel from our VCF (with `new_vcf2aHMM.py` ; see DRYAD/github repository), we set mean recombination rate to 0.7, a minimum rate of genotype calls for each parent population to three, minimum mean depth for the admixed population to 0.5 (1 read per individual), distance between sites to 5000 bp, and minimum allele frequency difference between populations to 0.7. We ran aHMM with a uniform 0.02 error rate and a single bidirectional hybridization pulse (50% from each population) at time 0.0001 to account for recent hybridization by disfavoring very small ancestry blocks. We validated parameters by running aHMM on known F1 hybrids and confirming heterozygous calls.

#### Ancient introgression

To determine whether *C. allenii* and *C. villosissimus* experienced ancient introgression in addition to the observed recent hybridization, we removed all individuals appearing as hybrids in the genetic PCA of wild individuals, leaving 28 *C. allenii* and 18 *C. villosissimus* individuals, and conducted ABBA-BABA tests for ancient gene flow (Graham and Farnsworth 2010; Durand *et al*. 2011; Soraggi *et al*. 2018). We tested for introgression across the whole genome and each chromosome with our population genetic samples using the -doabbababa2 1 function from the ANGSD package, specifying genome blocks of 5 Mb, to only include properly paired reads, and a minimum base and mapping quality of 30 (Soraggi *et al*. 2018). For ABBA-BABA test ingroup, we used a set of 21 *C. bracteatus* individuals that were collected, extracted, sequenced, and processed together with all the population genetic samples. The outgroup was the reference *C. lasius* genome. To calculate the D-statistic, standard error, and Z-score, we used ANGSD’s R script, `estAvgError.R` . For per-chromosome p-values, we used a false discovery rate (FDR) p-value adjustment to account for multiple testing across chromosomes (Benjamini and Hochberg 1995).

To evaluate whether evidence of introgression might be limited to Panama, we ran another ABBA-BABA test using a different dataset with a *C. villosissimus* individual from Ecuador, which is outside *C. allenii*’s range. For this analysis, we used capture sequencing data from 756 genes from Vargas *et al*. (2020). Because this dataset contains hard genotype calls, we used Dtrios from Dsuite to run the ABBA-BABA test (Malinsky *et al*. 2021). We included one *C. bracteatus* from Costa Rica as the ingroup, one *C. allenii* from Panama, one *C. villosissimus* from Ecuador, and two *C. lasius* from Panama and Peru as the outgroup (Figure 2B). The input VCF contained 25,978 variants that were divided into 20 Jackknife blocks (block size = 1297 variants) for calculation of the D-statistic.

#### Genome-wide differentiation

We calculated multiple differentiation and introgression statistics across the genome to characterize patterns and relate them to the genetic basis of habitat isolation trait between *C. allenii* and *C. villosissimus*. To characterize genomic differentiation, we generated all sites VCFs (including invariant sites) with bcftools mpileup, applied the same base filters described above in *DNA extraction, sequencing, and initial data cleaning*, and also removed sites with more than 80% missingness in either species. We then calculated the fixation index (*F*_*st*_; Weir and Cockerham (1984)), average nucleotide diversity within species (*π*), and average nucleotide difference between species (*D*_*xy*_) in 10 kb non-overlapping windows across the genome using PIXY (Korunes and Samuk 2021; Weir and Cockerham 2024). We also calculated the genome-wide *F*_*st*_ from 50 kb non-overlapping windows, summing variance components across all windows. To characterize introgression, we used a modified D-statistic ( *f*_*dM*_) that is better for sliding windows and depends on the derived allele frequencies across species (Malinsky *et al*. 2015; Martin *et al*. 2015; Malinsky *et al*. 2021). We calculated *f*_*dM*_ (and *d* _*f*_) in 50 variant windows with 25 variant steps with Dsuite’s Dinvestigate from VCFs filtered as above for quality, coverage and mappability (Malinsky *et al*. 2018b, 2021). We used two low-coverage *C. lasius* individuals and the *C. lasius* reference genome for the outgroup and 21 low-coverage *C. bracteatus* individuals for the ingroup, along with 21 low-coverage *C. allenii* and 18 low-coverage *C. villosissimus* genomes. We excluded all recent hybrids.

### Quantitative trait mapping

#### Trait measurement

The primary abiotic factors driving divergent adaptation are soil moisture and light (Chen and Schemske 2015). *Costus villosissimus* has several adaptations for seasonal drought that *C. allenii* lacks, and which cause mortality of *C. allenii* in *C. villosissimus*’ habitat (Chen and Schemske 2019). The first of these adaptations is seed dormancy: *C. villosissimus* seeds are dormant in the dry season whereas *C. allenii* seeds germinate as soon as they reach the soil. Adult *C. villosissimus* is adapted to drought through flexible drought deciduousness; during drought the plants limit water loss through active senescence of leaves (reabsorbing nutrients as leaves die). When rain returns, the plants take advantage of a higher light habitat to rapidly grow new leaves (Harenčár *et al*. 2022). Conversely, *C. allenii* is adapted to perennially wet, darker forest understory habitat and dies without substantial leaf nutrient reabsorption during drought. *Costus allenii* is also unable to take advantage of higher light to rapidly grow new leaves like *C. villosissimus* (Figures 1 and S6).

Differences in the biotic habitat select for *C. allenii*’s lower investment in growth as it trades off with higher investment in defense against herbivores. *Costus allenii*’s darker, wetter habitat holds a higher abundance of specialist herbivores. *Costus allenii* has thicker, tougher leaves with greater chemical defense against herbivores. These differences in defense between *C. allenii* and *C. villosissimus* result in specialist herbivores preferring to eat *C. villosissimus* in controlled trials, indicating that they would encounter higher levels of herbivory in *C. allenii* habitat (Harenčár *et al*. 2024).

##### Seed dormancy

Seed dormancy is an important adaptation in *C. villosissimus* to avoid drought, whereas *C. allenii* has no seed dormancy and seedlings die during drought in *C. villosissimus* habitat (Chen 2011; Chen and Schemske 2015, 2019). To score dormancy in our F3 mapping population, we tracked germination under controlled conditions in Percival seed incubators. Incubators were set to a 12-hour day length with a daytime temperature of 30°C and a nighttime temperature of 24°C. We collected and washed seeds, then placed them on consistently wet cotton. We recorded seed cohort - the batch of seeds harvested and sowed together. Twice a week, we surveyed all seeds for germination, which was determined by radicle emergence. We recorded the number of days from harvest/sowing in the incubators to germination as seed dormancy. After cotyledon emergence, we planted seedlings into four-inch pots in Pro mix HP+mycorrhizae and Biofungicide. We note that by quantifying dormancy in the seeds of the F3s we eventually genotyped, we avoid the problems associated with using dormancy values from the offspring of the genotyped individual; while we do not know whether the identified QTL impact seed dormancy through the seed coat, endosperm, or embryo, we do know they directly impacted dormancy in the genotyped individuals.

##### Growth rate

Rapid growth in *C. villosissimus* takes advantage of a higher light environment and enables drought adaptation through flexible drought deciduousness (Harenčár *et al*. 2022), whereas slower growth in *C. allenii* trades off with greater defense against herbivores in a lower light, higher herbivore pressure habitat. To measure growth rate in the F3 mapping population, we took height measurements when seedlings had at least 3-5 open leaves and again after 3-4 weeks. All seedlings were grown in growth chambers with 12-hour day length and 25°C constant temperature and were fertilized twice (start of week 1 and week 3) by watering evenly with 1 tsp Peter’s Professional NPK 20-20-20 fertilizer in 1 gallon of water. We calculated growth rate in cm per day as the total growth across all stems divided by the number of days between measurements. We tracked plant cohort (plants that reached 3-5 leaves and started in the growth chamber together) for use as an environmental variability covariate in trait mapping (see “Genetic map construction”). Within ∼ 1 week of the second growth rate measure, we moved plants from the growth chamber into the greenhouse and from half-quart pots to one-gallon pots.

##### Leaf nutrient reabsorption during drought

While *C. allenii* cannot survive without perennial water availability, adult *C. villosissimus* is adapted to drought through flexible drought deciduousness; during drought the plants limit water loss through active senescence of leaves (reabsorbing leaf nutrients like nitrogen and pigments like chlorophyll as leaves die). Conversely, *C. allenii*’s leaves simply wither and die without reabsorbing substantial nitrogen or chlorophyll (Figure S6). We quantified nitrogen and chlorophyll reabsorption during senescence in the mapping population as a proxy for drought deciduousness. We measured both leaf chlorophyll and nitrogen content before and after drought-induced leaf death. Mapping population hybrids grew for eight weeks in 1-gallon pots before we began the drought treatment. On the first day of no water, we recorded the average of three Soil Plant Analysis Development (SPAD; Minolta Camera Co., Osaka, Japan) meter readings of chlorophyll content taken near the tip, middle, and base of the oldest healthy leaf blade. From the same leaf, we also collected six hole punches of leaf tissue from the same locations as the SPAD readings, but on both halves of the leaf (one hole punch on each half of the leaf at the tip, middle, and base). We repeated the chlorophyll content measures and leaf hole punches when the leaf and the younger leaf above it were both fully dead. We dried all leaf tissue immediately after collection at 60°C for 72 hours and stored and shipped in silica. Leaf nitrogen concentration was determined with an elemental analyzer at Chapman University and at the UC Davis Stable Isotope facility. We quantified chlorophyll and nitrogen reabsorption as:

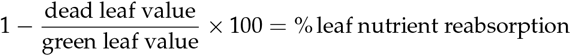

##### Leaf thickness and toughness

Leaf toughness and thickness are associated with herbivore defense and resource use strategy. To measure leaf thickness, we used a digital micrometer (Mitutoyo IP65, Global Industrial, NY, USA) to take measurements at three points on the 4th fully expanded leaf from the top of the stem. We took the average of measurements from the center of the leaf ca. 1 cm from the leaf edge, midway between the edge of the leaf and the midvein, and ca. 1 cm from the midvein. To measure leaf toughness, we used a Medio-Line 40300, 300 g max penetrometer (Pesola AG, Switzerland), to measure the force (g) required to puncture a 1 mm diameter hole in the leaf. We placed each leaf between two sheets of plexiglass with a 5 mm hole through both and centered the penetrometer rod in the hole, maintaining the leaf at a constant tension. We took the average of toughness measures collected at roughly the same three points as measured for thickness, transecting the middle of the leaf.

#### Ancestry assignment across hybrid genomes

We mapped local ancestry across the genome for each individual in the F3 mapping population using Ancestry HMM (Corbett-Detig and Nielsen 2017) and following the same methods as for the wild hybrids (see *Ancestry assignment across hybrid genomes* section above.

#### Linkage mapping

To identify and account for large structural variants and to get marker distance estimates in centimorgans (cMs), we generated linkage maps with Lep-MAP3 (Rastas 2017) using sequence data from the original wild F1, 21 F2s generated by selfing the F1, and the 390 F3s generated by selfing five of the F2s. We converted the genotype probabilities calculated with aHMM (above) into hard calls, keeping only sites with a call of greater than 90% probability. We incorporated the pedigree into the call file with `ParentCall2` . We used `SeparateChromosomes2` to assign markers to linkage groups, specifying a LOD limit of 11 and a theta of 0.001, and setting ‘distortionLod’ to 11 to filter for segregation distortion. To order markers within linkage groups we used `OrderMarkers2`, then plotted the cM position vs the reference bp position. We found that chromosome four was split into three linkage groups, likely due to a lack of informative recombination in our mapping population in large regions near the centromere; we manually joined these linkage groups. We also found and removed two small translocations in centromere regions (on chromosomes two & five) that could not be confidently placed. These modifications left nine linkage groups corresponding to the nine chromosomes (Harenčár *et al*. 2023). We re-ran `OrderMarkers2` 100 times for each chromosome and selected the run with the highest likelihood to use for cM position estimates.

#### QTL mapping

To identify regions of the genome associated with traits important for habitat isolation between these species, we performed QTL mapping with R/qtl (Broman *et al*. 2003). We used the markers and genotype calls with cM position from Lep-MAP3 described above. We dropped one individual with genotypes at less than 80% of markers, and removed markers found in fewer than 80% of genotyped individuals. We then thinned uninformative markers with `findDupMarkers`, leaving 14,889 markers with an average of 97.5% of individuals genotyped at any given marker. We also identified three pairs of individuals identical at almost all markers and removed one per pair. This left a total of 385 individuals with genotype data. We calculated genotype probabilities with the `calc.genoprob` function, and the recombination frequencies with `est.rf` . While most phenotype data was roughly normally distributed, seed dormancy had a long tail generated by unusually long dormancy times; we log-transformed seed dormancy to improve normality before model selection and fitting.

To determine appropriate covariates to include in the models for associating genotypes and phenotypes, we first used the R `step` function for stepwise selection via AIC, both through forward and backward elimination. We included hybrid index (percent *C. villosissimus* ancestry), F2 families, and cohorts as potential covariates. For each phenotype, we retained all covariates with a significant effect, using the ‘one-hot encoding’ method to recode cohort and family variables so they are treated categorically by R/qtl. This enabled us to avoid over-fitting by dropping from the models specific cohorts and families that were not significant covariates.

We then performed a genome-wide scan using a single QTL model with R/qtl’s `scanone` function with the Haley-Knott regression method (Haley and Knott 1992) to scan for associations between phenotypes and genotypes at ancestry-informative markers. We estimated and plotted 10% LOD (logarithm of odds) false discovery rate (FDR) thresholds for each phenotype with 1,000 permutations (S3). We used `fitqtl` to fit an initial model with the same covariates determined above and including the QTL peak or peaks that were above the FDR threshold.

To find the best model for all contributing QTL and their potential interactions, we ran `stepwiseqtl`, R/qtl’s automated function for forward/backward multiple QTL model selection, with penalties on main effects and interactions calculated with `calc.penalties` from 1000 permutations of `scantwo` for a 0.05 alpha. For computational tractability, we reduced the marker set with another, more stringent round of duplicate removal, and calculated genotype probabilities at 5 cM steps with an error probability of 0.01. We then ran `fitqtl` on the original, larger cross object with genotype probabilities calculated at marker locations using the model and QTL peak positions selected by `stepwiseqtl` after running `refineqtl` on the peak positions with the original cross object. To determine the size of the QTL windows, we used R/qtl’s `bayesint` function to calculate 95% Bayes credible intervals.

We also note that iterating `refineqtl` to refine QTL peak positions, `addqtl` to add additional peaks one at a time, and `fitqtl` to check for significance of the additional peaks identified multiple additional peaks for some phenotypes. Here, we conservatively only report results from the model identified via a penalized LOD score with `stepwiseqtl`, other than for growth rate and percent chlorophyll reabsorbed during drought. For growth rate, 1000 permutations of scanone identified a peak on chromosome 3 that was not included in the model from `stepwiseqtl`, but that is significant at a p-value of <0.01 when included in the model; we report results from this model. For percent chlorophyll reabsorbed during drought, `stepwiseqtl` erroneously identified two peaks within one broad peak on chromosome 1. The two loci identified have zero recombination events between them in our mapping population, and thus were part of one QTL; we report results from modeling a single QTL.

#### Relating introgression to the genetics of adaptive divergence

To investigate patterns of introgression and differentiation around the QTL loci that contribute to adaptive divergence, we compared genome-wide *F*_*st*_ to the average *F*_*st*_ within QTL windows. For each chromosome, we plotted the QTL peaks and credible intervals over *F*_*st*_ calculated in 10kb non-overlapping windows with 99th percentile outliers highlighted in red (Figures 3C and S4). Next, we calculated windowed genome-wide *F*_*st*_ averages and compared them with QTL window averages. We first estimated cM positions for the summary statistic windows by fitting isotonic regressions to each chromosome using the cM and bp marker positions from Lep-MAP3. We excluded chromosome 4 from this analysis due to poor genetic mapping. We then averaged *F*_*st*_ in one cM windows across the genome and for each QTL window that was less than 15 cM ( ∼ 10 Mbp). This excluded the windows for nitrogen reabsorption during drought on chromosome 2 and the for growth rate on chromosome 5, which had CIs larger than 15 cM. Finally, we compared the distribution of genome-wide windowed *F*_*st*_ values to QTL window averages (Figure 3D). We applied the same method to *D*_*xy*_ and *f*_*dM*_, with *D*_*xy*_ one cM window values averaged from 10 kb PIXY windows just like *F*_*st*_, and *f*_*dM*_ averaged from Dsuite 50-variant, 25-variant step sliding windows (Figure S5).

## Data, Materials, and Software Availability

Sequence data are available on NCBI SRA under BioProject PRJNA1479894. All other data and code are available on DRYAD (DOI: 10.5061/dryad.xgxd254xj).

## Acknowledgments

We gratefully acknowledge Dr. Jennifer Funk for coordinating leaf nitrogen quantification and Monica Nguyen and Madalyn Papenfuss for processing the leaf tissue. For critical plant care, greenhouse facility management, and excellent growing advice, we thank the UCSC greenhouse team, including Sylvie Childress, Jim Velzy, and Laura Palmer. For greenhouse plant care, trait measurement, data (double) entry, and DNA extraction, we enthusiastically thank Reed Hammock, Kelsey Cochran, Hannah Thacker, Wade Matern, Erin Thompson, Avery Hansen, Aubrie Tait, Melina Carrizosa, Madison Evanow, Abigail Holtz, Alyssa Kattman, Samson Janse, Janine Anne Tan, Adrian Flores, and Carolina Saucedo, without whom the genetic mapping would have been truly impossible. We express deep gratitude to Drs. Merly Escalona and Maximilian Genetti for computational assistance, Rion Parsons for computing cluster support, and the UCSC PaleoGenomics lab for input on population genomic analyses. We also thank Douglas Schemske, Sharon Strauss, Johanna Schmitt, Jeffrey Ross-Ibarra and his lab, Graham Coop and his lab, Jennifer Gremer and her lab, Beth Schapiro and her lab, Russ Corbett-Detig, Molly Schumer, and Ingrid Parker for feedback on lab methods, analyses, figures, and/or the manuscript. The sequencing was carried out at the DNA Technologies and Expression Analysis Cores at the UC Davis Genome Center, supported by NIH Shared Instrumentation Grant 1S10OD010786-01. Research along Pipeline Road in Panamá was conducted under Permiso Especial SE/P-13-19 from Autoridad Nacional del Ambiente and Ministerio de Desarollo Agropecuario del República de Panamá no. autorización 29363 and MiAmbiente permit ARB-022-2022. Generative AI was not used int he writing of this manuscript, but was used for code optimization, some data visualization code generation, and data and code organization and reproducibility testing. This research was funded by the Jean H. Langenheim Endowed Graduate Fellowship, the Jean H. Langenheim Chair in Plant Ecology and Evolution, the Achievement Rewards for College Scientists Scholar Award, the UCSC Plant Science Fund for greenhouse charges, a Tinker Foundation Field Research Grant, the P.E.O International Scholar Award, the Garden Club of America Fellowship in Tropical Botany, the American Society of Naturalists Student Research Award, the UCSC Ecology and Evolutionary Biology department’s summer funding for graduate students, and National Science Foundation Dimensions of Biodiversity grant DEB 1737889 to K.M.K.

## Supplement

### Supplemental figures

**Figure S1.**
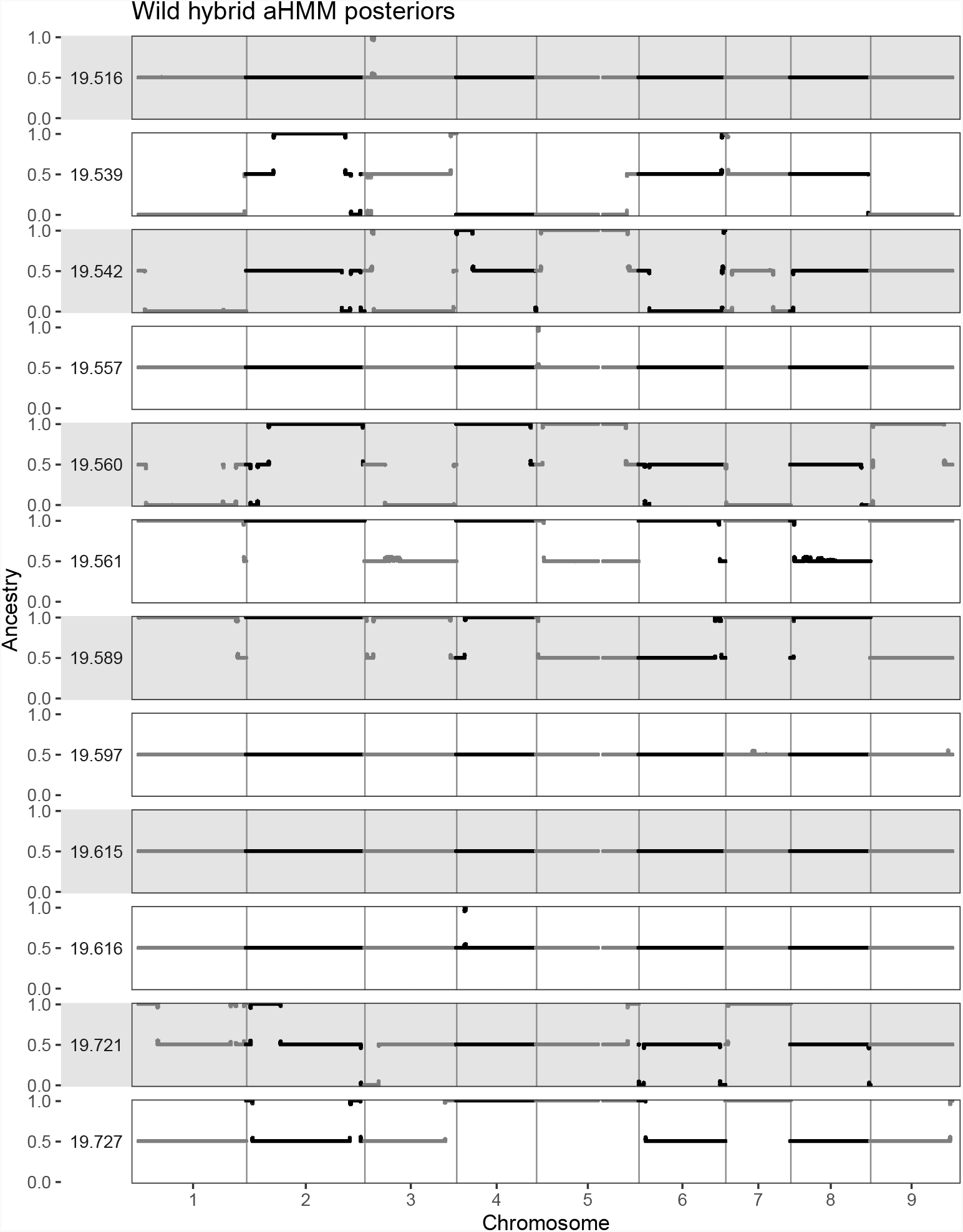
Local ancestry inference plots for wild Panamanian hybrids. The x-axis is position along the chromosome and the yaxis represents ancestry at that position, with 0 indicating homozygous *C. allenii* ancestry, 0.5 indicating heterozygous ancestry, and 1 indicating homozygous *C. villosissimus* ancestry.

**Figure S2.**
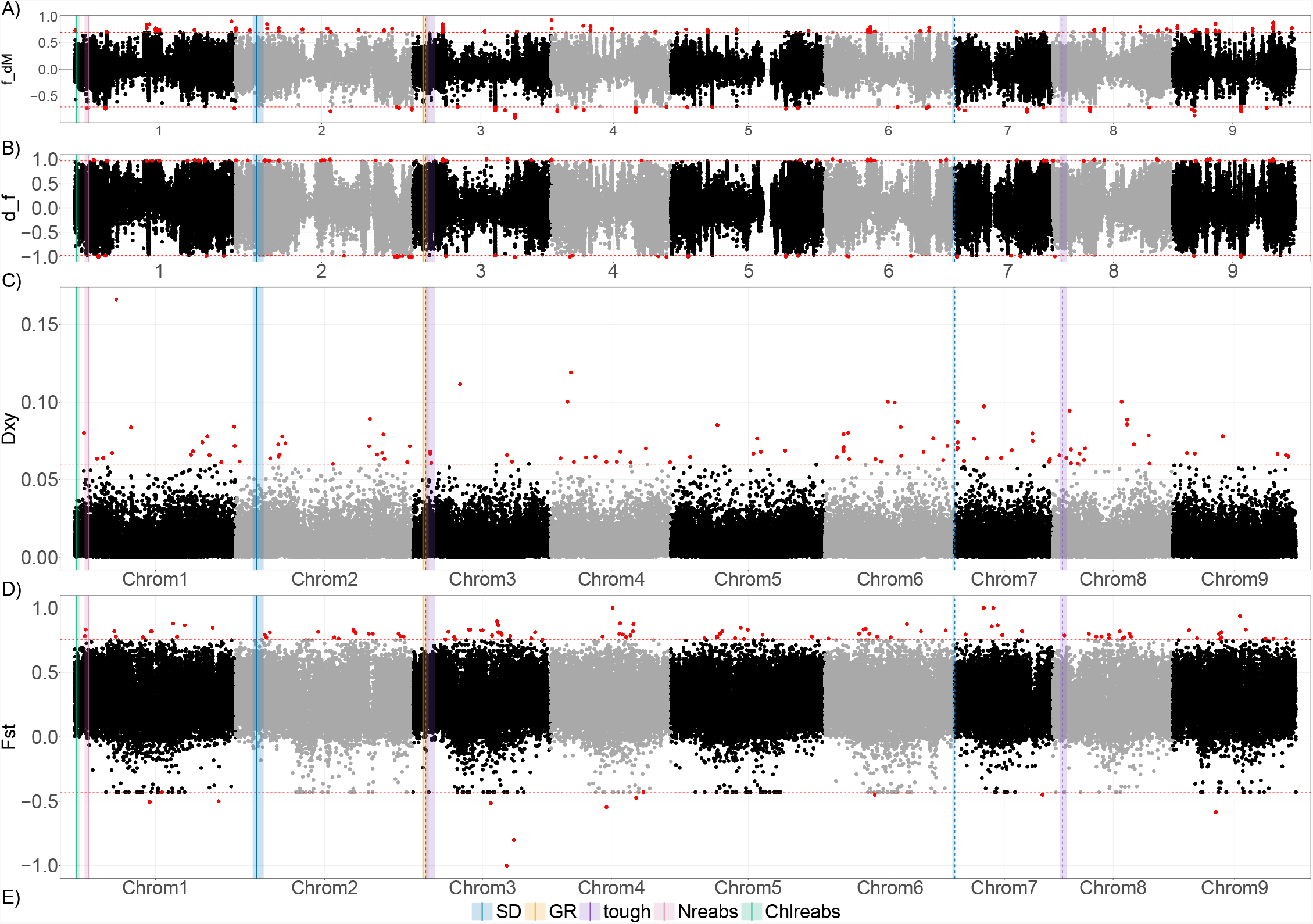
Genomic patterns of introgression and differentiation. A) *f*_*dM*_, B)*d*_*f*_, C) *D*_*xy*_, D)*F*_*st*_. For both introgression statistics, *f*_*dM*_ and *d* _*f*_, positive values indicate a signal of introgression between *C. allenii* and C. villosissimus, values near 0 indicate no introgression, and negative values indicate a signal of introgression between *C. bracteatus* and *C. villosissimus*. Red lines represent the 99.9th percentile of negative values, and red dots represent points either below this value, or above its absolute value. Points above the absolute value indicate loci with evidence of introgression between *C. allenii* and C. villosissimus. Both *d* _*f*_ and *f*_*dM*_ were calculated in 50 variant sliding windows of 25 variant steps. Genomic patterns of diversity and differentiation (*D*_*xy*_ and *F*_*st*_) were calculated in 10kb non-overlapping windows. Colored bars indicate QTL peaks with dashed bars indicating antagonistic QTL and shaded regions showing Bayes 95% CIs.

**Figure S3.**
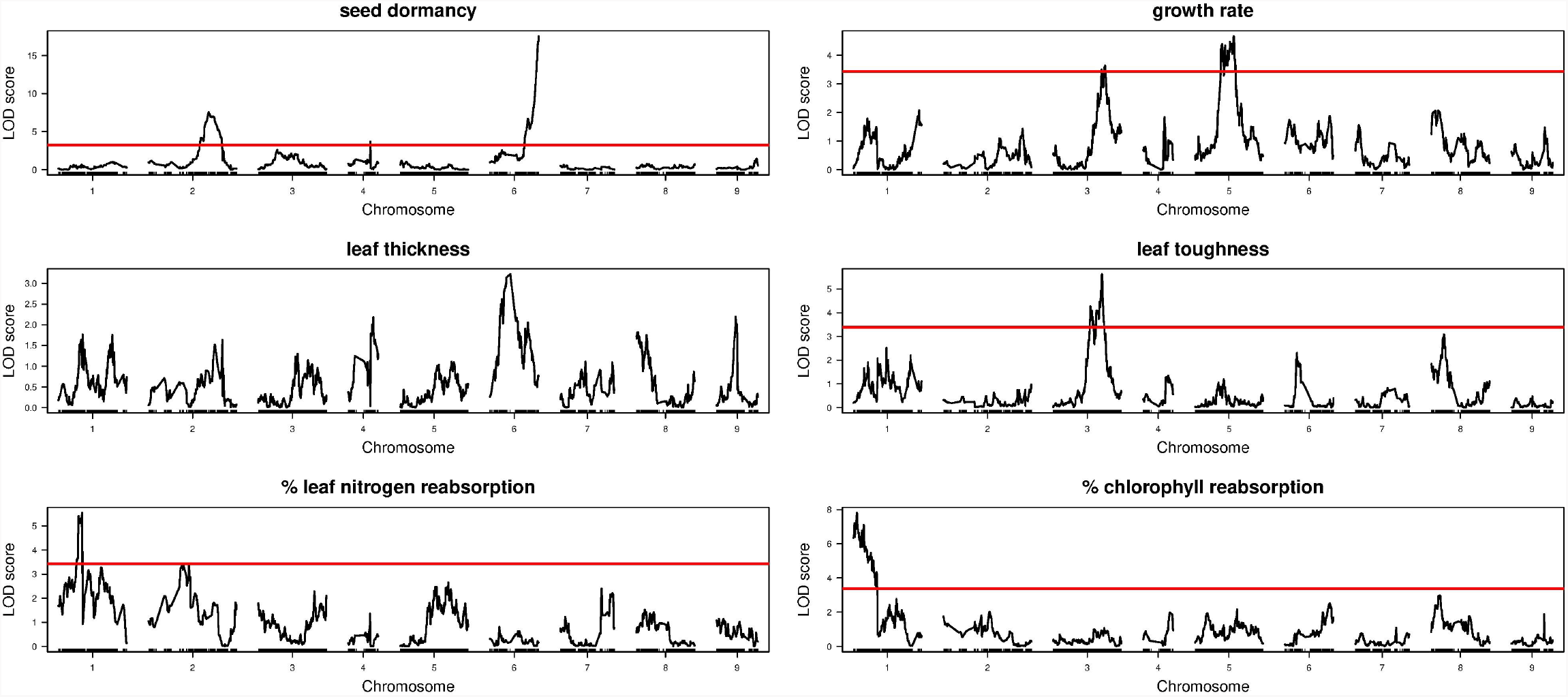
QTL peaks with 10% LOD score cutoffs in red calculated from 1000 permutations of `scanone` .

**Figure S4.**
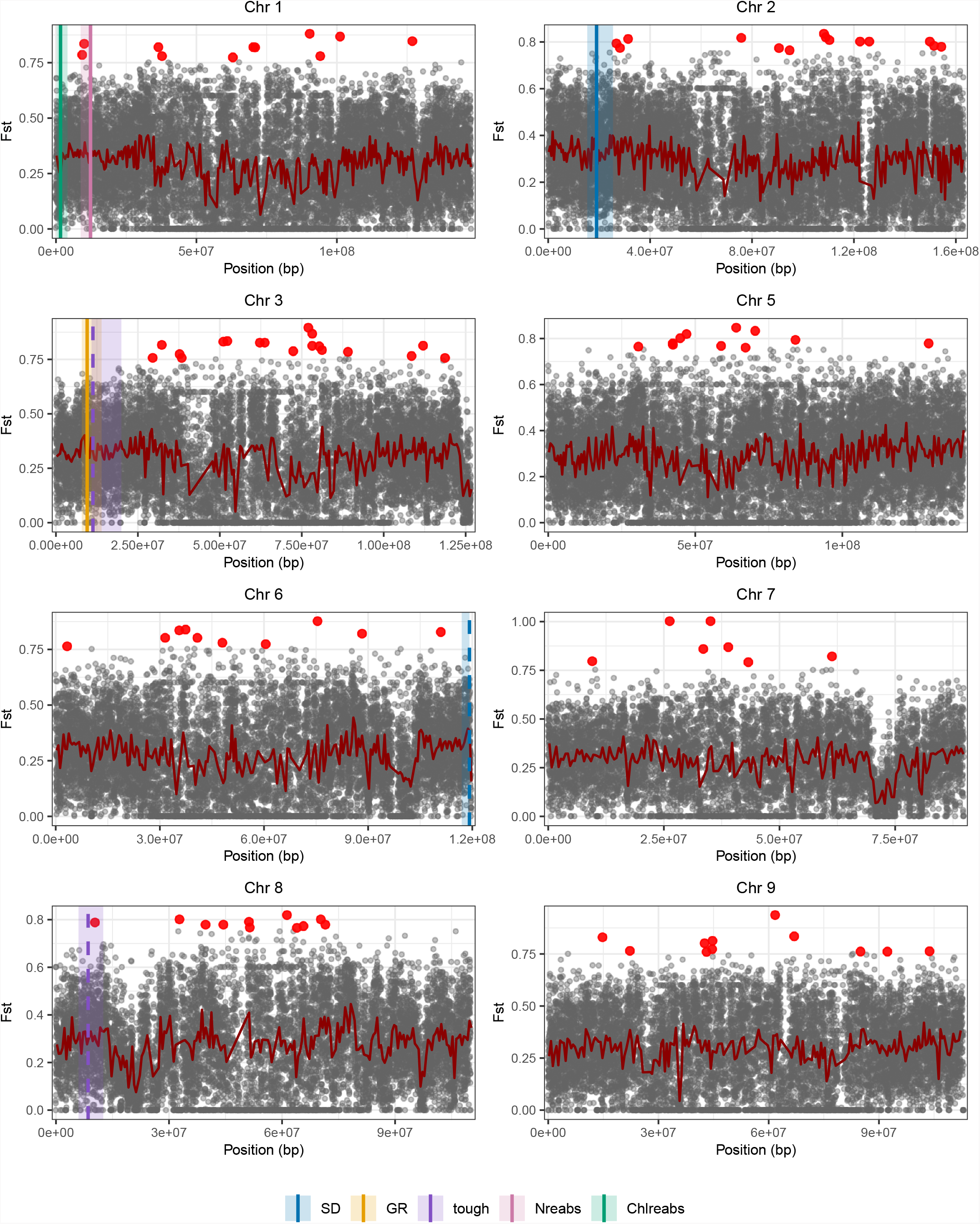
Genomic patterns of differentiation with QTL. Genome-wide *F*_*st*_ calculated in 10 kb windows with overlaid QTL. Red points are 10 kb windows in the 99.9th percentile and the red line indicates *F*_*st*_ average calculated in 500 kb windows. Colored bars indicate QTL peaks with dashed bars indicating the antagonistic QTL and shaded regions showing Bayes 95% CIs.

**Figure S5.**
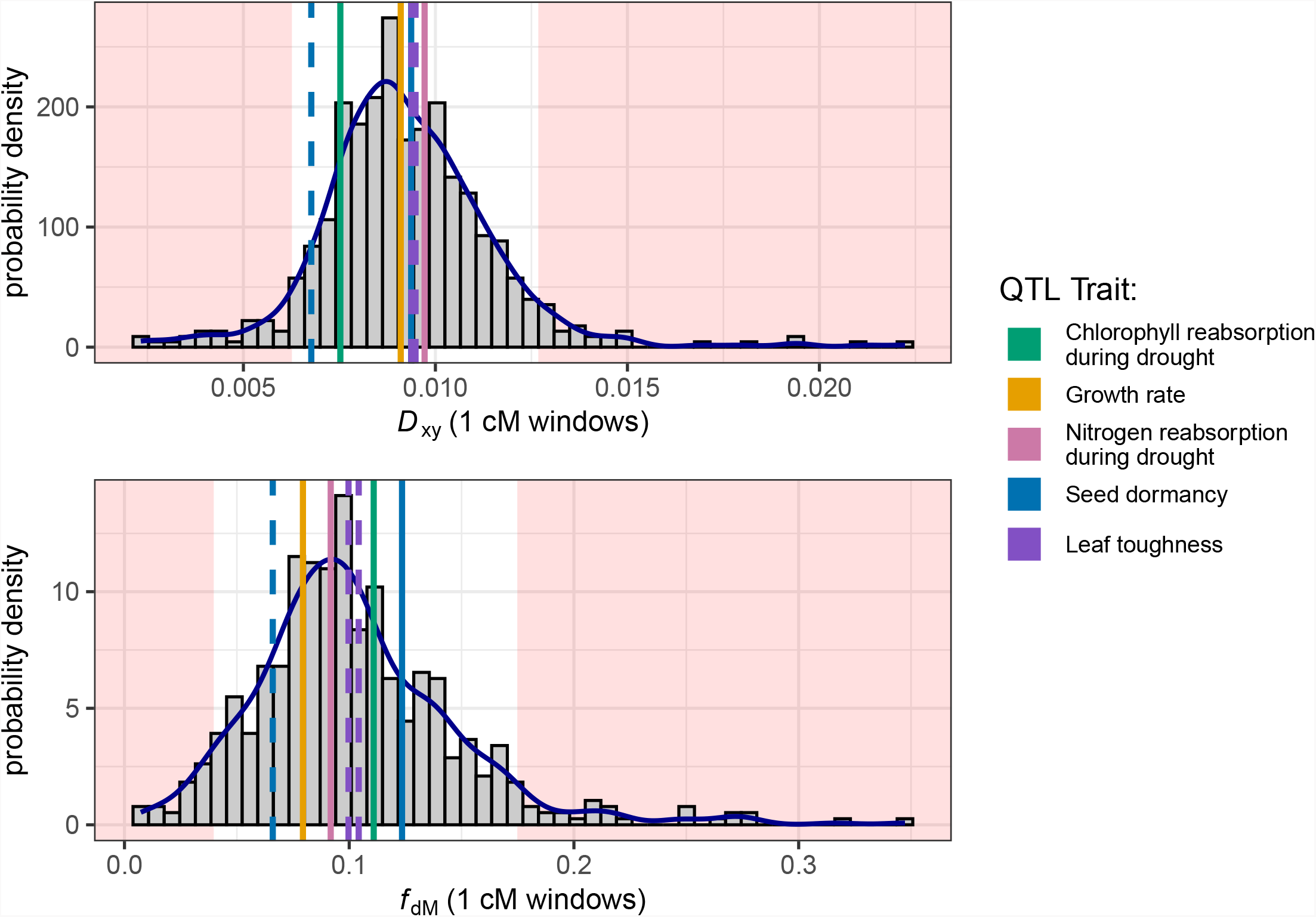
QTL-window average *D*_*xy*_ and *f*_*dM*_ relative to genome-wide distributions. QTL-window *D*_*xy*_ and *f*_*dM*_ compared to genome-wide one cM window averages with 5th and 95th percentiles shaded red. (A) *D*_*xy*_ from PIXY (10 kb windows). (B) *f*_*dM*_ from Dsuite Dinvestigate (50-variant windows, 25-variant step). See Table S2 for QTL intervals.

**Figure S6.**
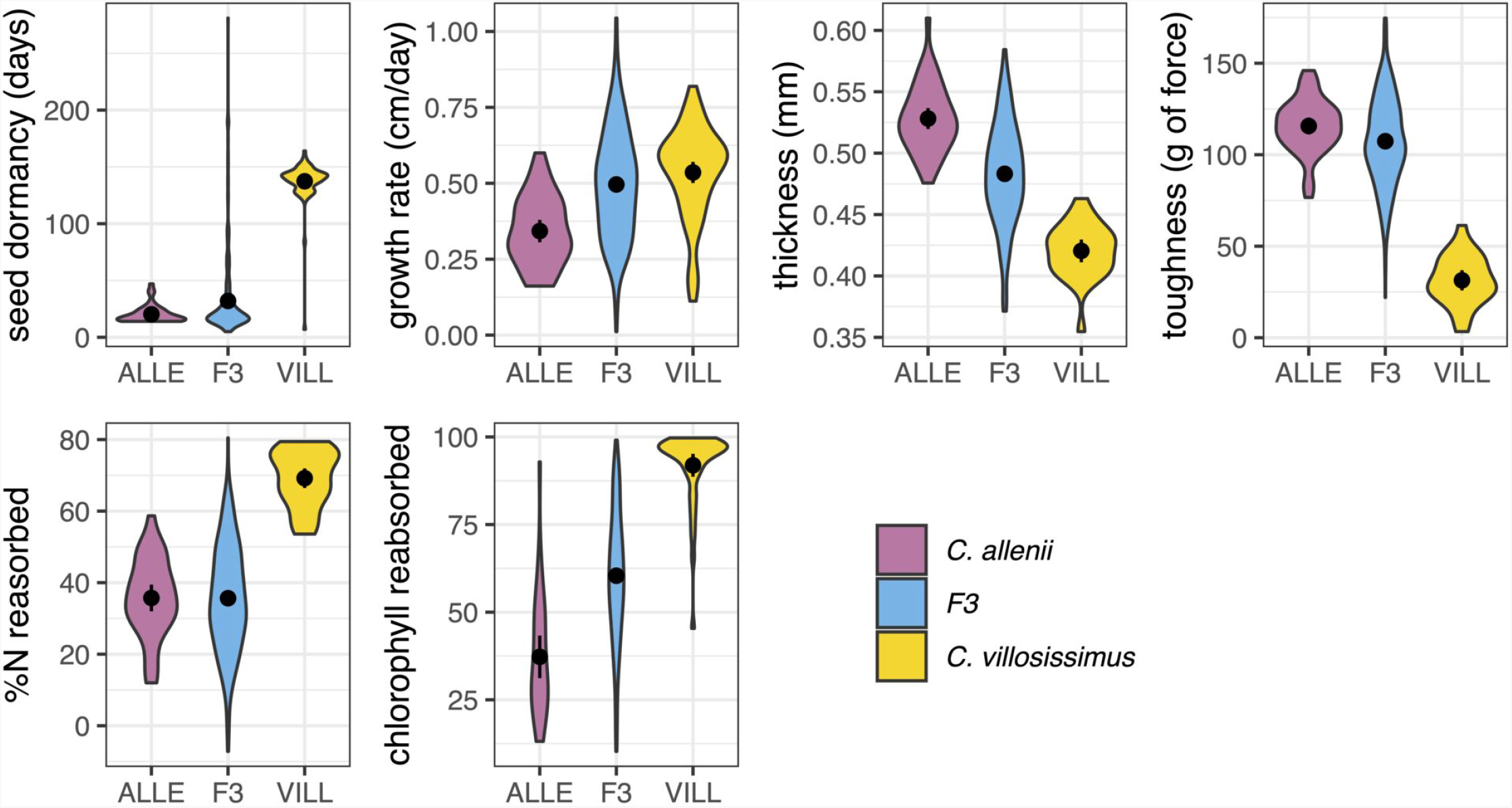
Trait distributions for the F3 mapping population and parentals. All trait data are from plants grown in the same growth chambers (early life stages) and greenhouse.

### Supplemental tables

**Table S1.**
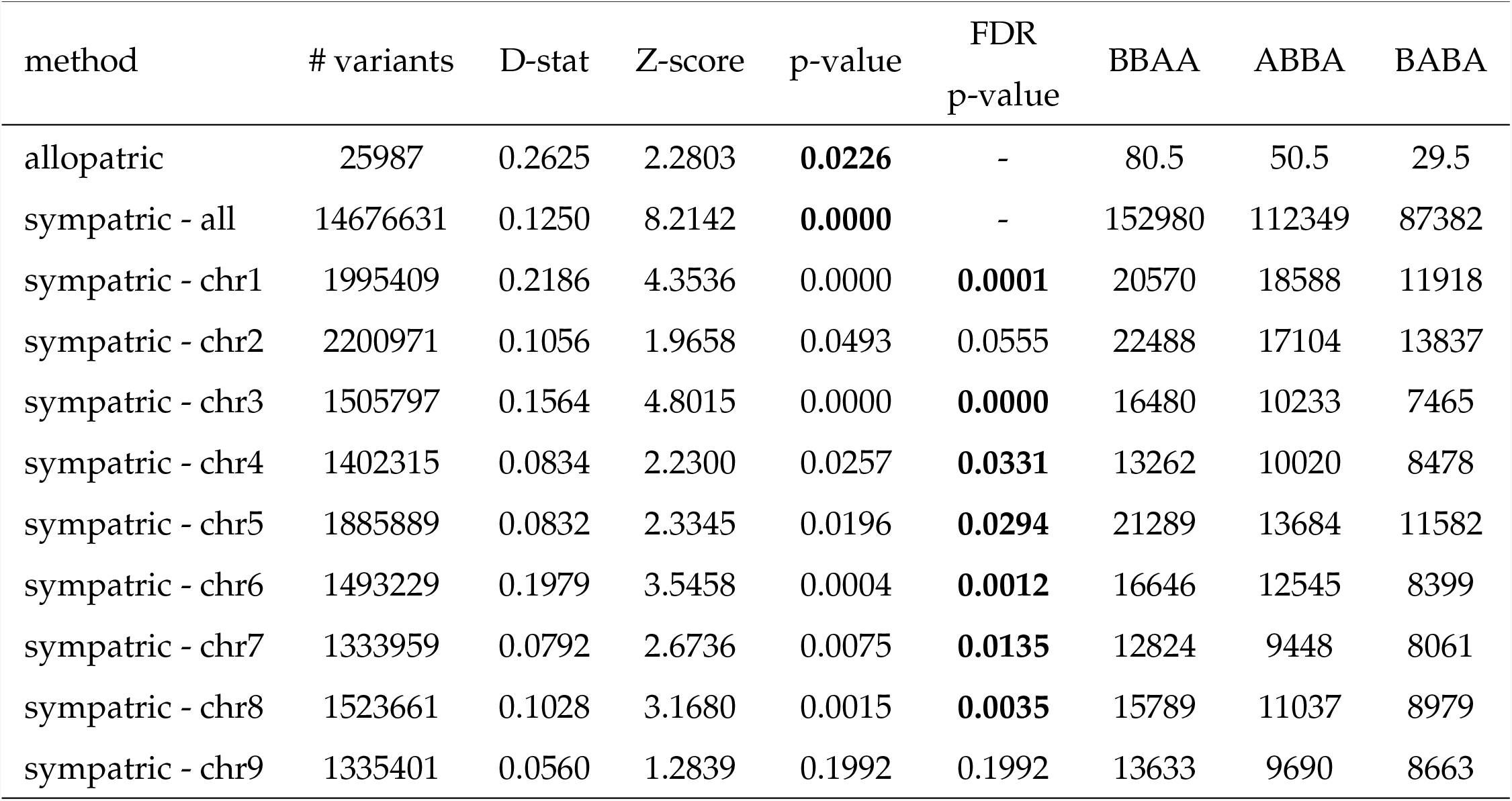
Results from ABBA-BABA tests run on population genetic samples with angsd -doAbbababa2, and on capture-sequencing data with only a non-Panamanian *C. villosissimus* using Dsuite’s Dtrios. In all cases, P1 = *C. bracteatus*, P2 = *C. allenii*, P3 = *C. villosissimus*, and Outgroup = *C. lasius*. For repeated tests on each chromosome, we applied Benjamini and Hochberg (1995) p-value adjustments to account for multiple testing.

**Table S2.**
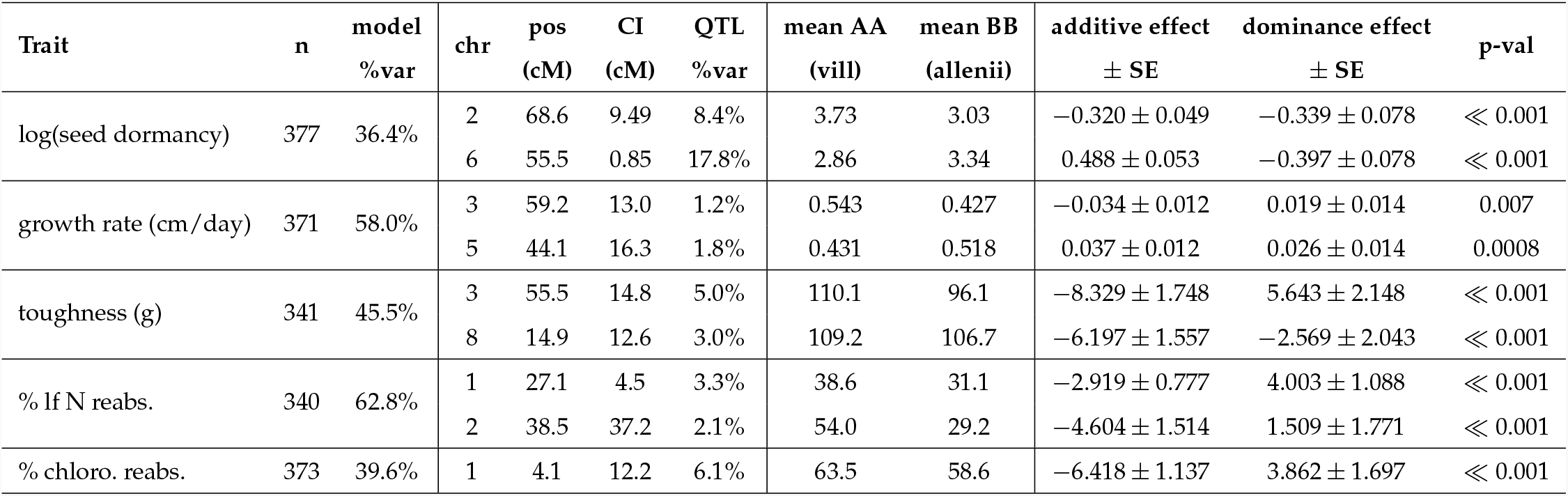
QTL locations and effect statistics. Columns: trait, number of individuals phenotyped, total variance explained by the model with the listed QTL peaks and covariates, chromosome, position of the highest LOD score (cM), 5% Bayes credible interval width (cM), QTL %variance, additive and dominance effects *±* SE, and QTL p-value. raw genotypic means at the peak marker for AA (*C. villosissimus*) and BB (*C. allenii*); additive and dominance effects *±* SE from `fitqtl` ; and QTL *p*-value. Additive effects are *a* = (*µ*_AA_− *µ*_BB_)/2 after covariates. Dominance is *d* = *µ*_AB_ − mid-parent, or the difference between the heterozygote and the midpoint between homozygotes. For nitrogen and chlorophyll reabsorption, raw means show AA > BB (matching higher reabsorption in *C. villosissimus*), but the covariate-adjusted additive effects are negative: after accounting for background genetic effects, residual association at the locus flips relative to the raw genotypic means.

